# In-depth characterization of Staurosporine induced proteome thermal stability changes

**DOI:** 10.1101/2020.03.13.990606

**Authors:** Alexey L. Chernobrovkin, Johan Lengqvist, Cindy Caceres Körner, Daniele Amadio, Tomas Friman, Daniel Martinez Molina

## Abstract

Cellular thermal shift assay (CETSA) coupled with high-resolution mass-spectrometry (MS) has proven to be indispensable tool to track thermal stability changes in cellular proteins caused by various external or internal perturbations. One of the major applications of CETSA MS is still binding profile characterization of small molecule drugs. Before applying the method for characterization of novel compound it is crucially important to understand the limitations, sensitivity and ways to improve the throughput of the different method implementations. Here we present deep comprehensive profiling of Staurosporine induced proteome thermal stability alteration utilizing different experiment layouts. By applying unbiased straightforward sample preparation and data analysis approaches we were able to compare and benchmark different experiment layouts for detecting compound-induced protein stability changes in CETSA MS experiments.

## Introduction

In drug discovery, achieving high efficacy and safety requires a deep understanding of both the mechanism of action of the pharmacologically active compound and the therapeutic target(s). To develop the most effective drug, the researchers/developers must determine the role of the target in the disease of interest, as well as how it interacts with and is modulated by the compound. However, even if the primary target is known, it is crucial to make sure a full understanding of the complexity of the mechanism of action (MoA), including the effect on the primary target and any off targets. In this regard, proteome-wide, mass spectrometry (MS)-based methods have emerged as powerful tools. MS-based proteomics typically involves two main approaches: chemoproteomics, where either target protein or ligand is being chemically labelled, label-free approaches, where changes in physico-chemical properties of either protein or ligand are measured. While each method has its advantages, it is valuable to consider which approach will be most vital to the research success in the long run. Cellular thermal shift assay (CETSA) was introduced in 2013^1^ and soon proved to be an indispensable tool to measure and validate target engagement in such complex samples as living cells or even tissues. Following the proof-of-concept paper, CETSA was combined with mass-spectrometry based quantitative proteomics (CETSA MS or TPP)^2^ which allowed to profile changes in protein thermal stability on the proteome-wide basis.^3–6^ In the first implementation, aliquots of sample with or without compound were heated to the range of temperatures, and melt curves for thousands of proteins were measured simultaneously using isobaric labeling quantitative proteomics.^2^ Changes in thermal stability were assessed by measuring the difference in T_m_ values between compound-treated sample and vehicle control. Furthermore, by measuring isothermal concentration response at a constant heating temperature (usually around a protein’s melting temperature) for the specific protein of interest one can not only *identify* protein-ligand binding, but also rank compounds according to the binding affinities by measuring isothermal concentration response at the constant heating temperature (usually around protein’s melting temperature). However, to explore concentration domain not for a single protein but for the whole proteome one need to be able to measure concentration-response curves at a number of different temperatures covering the whole proteome melting range. This approach was implemented in 2D thermal proteome profiling (2D-TPP)^4^ and later also in the so-called IMPRINTS format^3,6^ where fewer temperatures were analyzed. These approaches were successfully applied to profile cell-cycle driven changes in protein stabilities.^6^ In order to assess thermal stability changes of a specific protein or class of proteins it is even possible to perform proteome wide analysis at a single temperature point, but this however would discriminate proteins melting at much higher/lower temperatures. Alternatively, instead of single-temperature measurements, full melt curves or the 2D-TPP format, one can integrate (compress) melt curves in vitro and experimentally assess the difference in areas under the melting curves at a predefined range of temperatures.^7,8^ In such case, decreased due to the melt curve compression sensitivity can be well compensated by substantial increase in throughput and number of possible replicates. In this manuscript we assess the pros and cons of different acquisition strategies for measuring protein stability alteration. We chose Staurosporine, a pan-kinase inhibitor, that binds to a large number of proteins (mainly kinases), resulting in both stabilization and destabilization, to different extents and at protein specific affinities, thus giving us the possibility to correlate the responses measured by different methods.^9–11^

## Results

In order to be able to compare different CETSA MS formats we’ve kept sample preparation, MS analysis and data analysis methods as unified as it possible (see details in methods). In all experiments we’ve characterized protein hits (e.g. proteins changing stability upon Staurosporine treatment) by effect size (amplitude) and significance (−log10(p-value)).

To make effect sizes comparable across different experimental layouts we’ve used log2-transformed ratio of areas under the melt curves (AUMC) instead of the commonly-used ΔT_m_ as a measure of stabilization in comparison of protein melt curves:

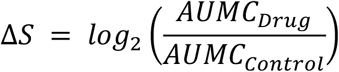

In the compressed format we directly derive ΔS as a log2-transormed fold change of soluble protein intensity relative to untreated control. As an effect size in such experiments we use the ΔS value corresponding to the highest compound concentration applied.

To assess significance, we’ve performed non-linear fit of temperature and/or concentration-dependent changes in soluble protein amount and compared it to the trivial (e.g. no stability change) model via ANOVA F-test. It allows for direct comparison of sensitivities and specificities of different acquisition formats regardless of the particular experiment layout.

### Melt curve experiment in lysed K562 cells

Lysed K562 cells were treated with the compound (20 μM of Staurosporine in 1% final DMSO), and with vehicle control in duplicates for 15 minutes at room temperature. The amount of proteins that remained soluble after 3 minuntes of heating to temperatures 40–67°C was measured using TMT-based quantitative proteomics. Experiment layout is very similar to what was reported in the original Savitski paper^2^, except for the slightly different temperature range. We have identified 8321 protein groups; for 6,351 protein groups it was possible to fit the melting curve in both treatment conditions.

Melt curve analysis allows us to identify 172 proteins changed their stability, with 96 proteins being stabilized and 76 destabilized by Staurosporine (Figure 2, Supplementary figure 1). Remarkably, substantial part of stabilized proteins were annotated as kinases - <u>68 proteins</u>, or 70%. Another 8 proteins are known binders of protein kinases. However, only <u>~15</u>of 76 destabilized proteins were annotated as kinases or ATP binders.

**Figure 1.**
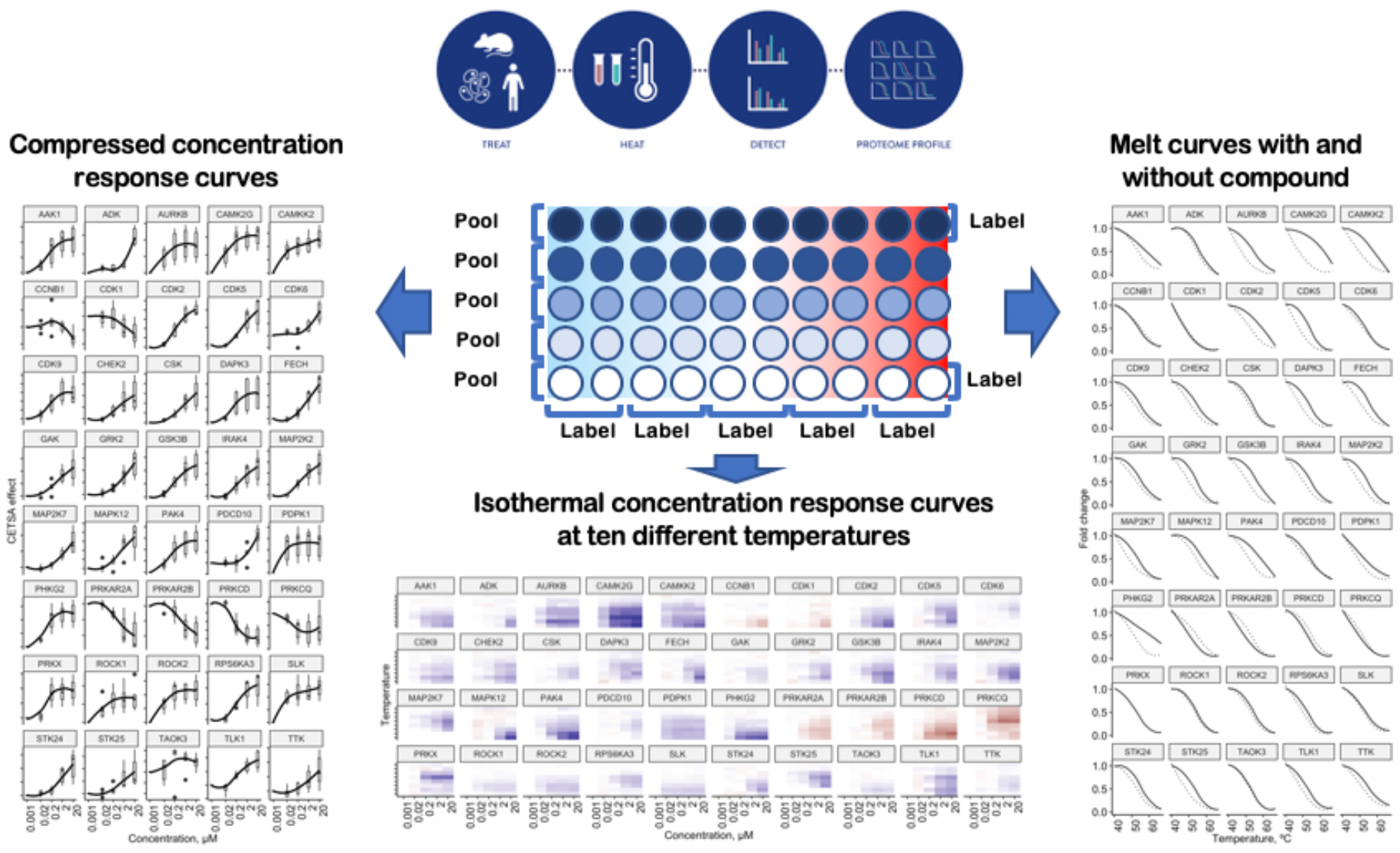
Schematic representation of the different experimental layouts to measure compound-induced changes in protein thermal stability. Measurement of soluble protein at different heating temperatures for single compound concentration corresponds to the melt curve format (right panel); different temperatures and different concentrations presented as heatmaps – 2D format (middle) and compressing melt curves at different compound concentrations corresponds to the compressed format (left panel).

**Figure 2.**
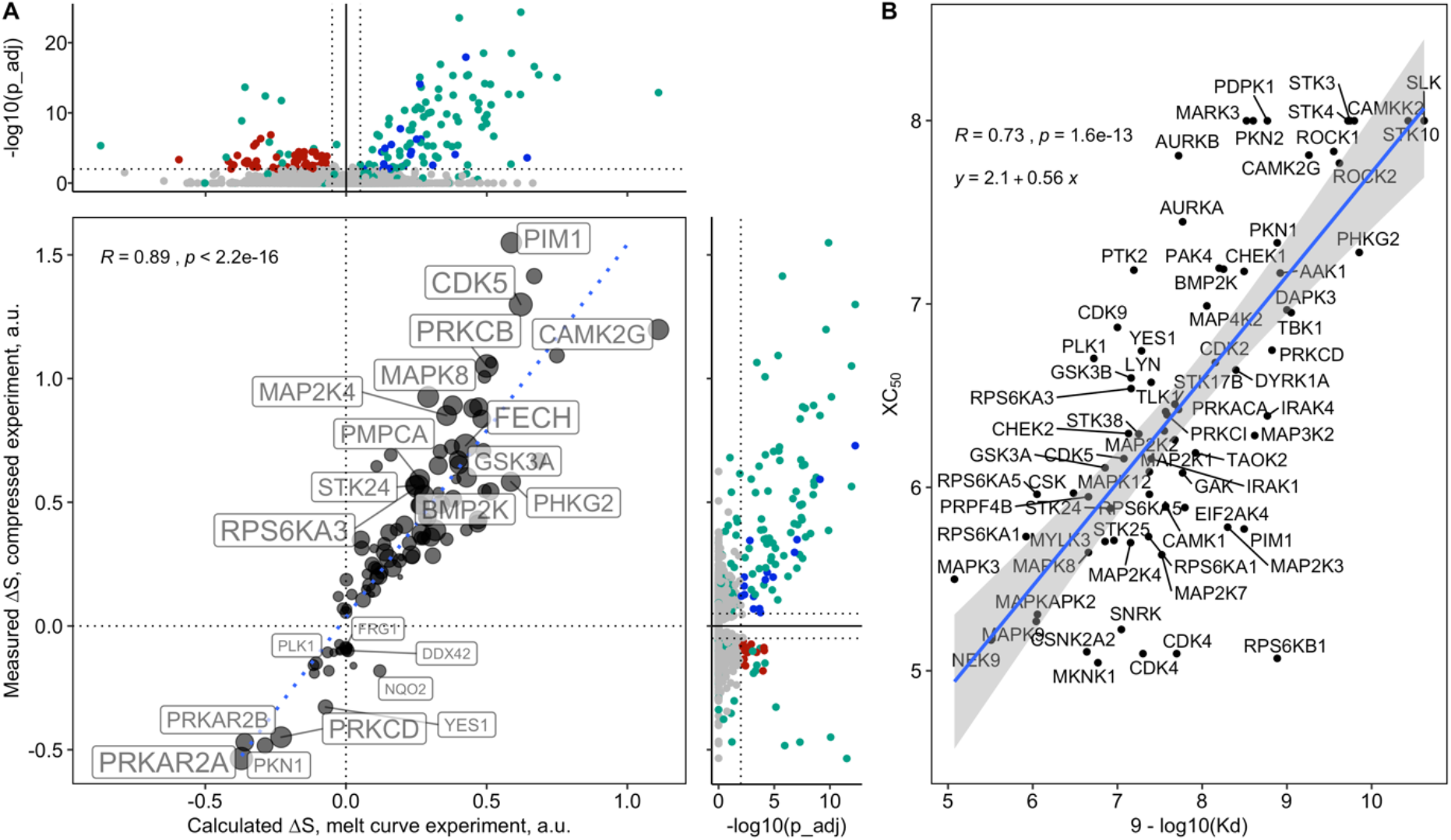
CETSA MS results of two experiments-melt curves and compressed. Panel A show the volcano plots were kinases and kinase-related proteins shown in green, other stabilized proteins in blue, destabilized in red, non-significant-in grey, and a scatter plot comparison of ΔS values in two experiments. Panel B shows the correlation between XC_50_ values from CETSA experiment and K_d_ values from competitive binding assay.

In order to study relation between ΔT_m_ values and ΔS values (Eq1), we’ve calculated them for all protein melting curves. Strikingly, ΔT_m_ and ΔS values show almost ideal linear correlation, with Pearson’s correlation coefficient of 0.97 (Supplementary figure 1B), suggesting that area-based ΔS values are at least as good as previously used melting temperature shifts). Worth to mention, that stability alterations as small as ΔT_m_ of 0.5 °C (ΔS~0.08) were detected at the selected significance level, and no artificial, manually defined, filtering criteria except for significance cut-off were used here, making suggested statistical approach more favorable compared with previously used approaches^2^.

### Compressed, 5 concentration points

As we have shown previously, melt curve format provides easy to understand results with a large number of protein kinase hits. However, we won’t get any idea about potencies, K_d_ or any other concentration-dependent differences in target protein engagement as only a single compound concentration is tested. However, 2D-TPP format^4^ allows to assess protein stabilizations at 5 concentration points, which is enough in some cases to estimate XC_50_ values and use them in the follow-up analysis. This exhaustive format comes with a cost – increased analysis time and much more complicated data analysis and interpretation. In addition, much higher number of samples to handle, which inevitably leads to systematic errors and trends in the data.

The “Compressed” format, or one-pot analysis, as has been suggested earlier, allows to significantly reduce the number of samples analyzed by integrating the protein melt-curve in vitro. To assess efficiency and limitations of such approach, and to compare it to the melt curve and 2D-TPP approaches, we’ve performed Compressed CETSA MS analysis with 5 concentration points (same as in 2D). In this case, the five concentration points were: Control (0), 0.02, 0.2, 2 and 20 μM. The soluble fractions from 10 different temperatures (40-67°C, same as in the MC experiment) were pooled together and proteome-wide concentration-dependent changes in protein thermal stabilities were measured using TMT-based quantitative proteomics. We have performed four biological replicates, which resulted in two TMT-10 labeled samples to analyze (compared to four in MC and five in 2D). A comparison of results between melt curves and compressed experiment results are provided in Figure 2, while the summary of the compressed experiment results are provided in the Supplementary Figure 2.

From the results it is clear that time less than it is required for MC or 2D experiment, we’ve been able to detect similar number of Staurosporine-induced changes in protein thermal stability: 83 proteins (74 kinases) were found to be stabilized and 17 proteins (15 kinases) destabilized upon treatment with Staurosporine. The measured compressed ΔS values correlate well with ΔS values measured in MC experiment (Figure2A, supplementary figure 3B), once again suggesting that different acquisition formats indeed measure the same thing at the end.

However, unlike the MC format, here we have also covered concentration domain, and thus can extract information about *target engagement potencies* for different protein targets of Staurosporine. For those proteins were concentration-response curve were fitted with reliable goodness R^2^>0.8, CETSA-derived XC_50_ may give us a good indication about ligand interaction with native protein in the very complex sample matrix. Figure 2B shows correlation plot of CETSA-derived XC_50_ and previously published K_d_ values measured in competition binding assay on synthetic kinases.^10^ Significant correlation of XC_50_ and K_d_ suggest the relevance of measured XC_50_ values.

### Compressed, 8 concentration points

After assessing the effect of Staurosporine in K562 cell lysates we decided to also investigate the effect in intact cells alongside the corresponding cell lysates. In the intact cell format, the cell physiology is maintained and therefore are more pathway effects are anticipated. For this study we also extended the concentration range to eight points in order to get a better curve fit. Moreover, non-heated controls were also included in order to control for non-CETSA effects, *i.e.* effects that are not due to changes in thermal stability. A broadly acting drug such as Staurosporine could induce changes in protein availability, which could stem from changes in protein degradation or synthesis. Also, altered subcellular localization could contribute to changes in protein availability.

The intact cell experiment resulted in more significantly shifted proteins (referred to as hits) compared to the lysate format; 485 vs. 83 at p<0.01 (Figure 3, Suppl. Tables). In the intact experiment, 67 kinases (52 being protein kinases) were identified as hits compared to the lysate format where 77 kinases (75 being protein kinases) were identified, and 34 protein kinases were overlapping between formats. Only four non-kinase proteins were overlapping between formats: Ferrochelatase, mitochondrial (FECH), Thioredoxin-dependent peroxide reductase, mitochondrial (PRDX3), mitochondrial peptidase subunits alpha (PMPCA) and beta (PMPCB). The majority of the hits in the intact cell experiment were non-kinases and many of these could be mapped to protein kinase signaling pathways by string analysis (Supplementary Figure 3), indicating that perturbation of kinase activity in intact cells also affected the thermal stability of kinase-interacting proteins. Indeed, Staurosporine treatment of intact cells had a significant effect on the thermal shifts of several non-kinase proteins belonging to multiple signaling pathways. Prominent examples of affected non-kinases include the GSK3B-interacting protein GSKIP and the cyclin dependent kinase substrate RB1 (Fig. 3 B, C).

**Figure 3.**
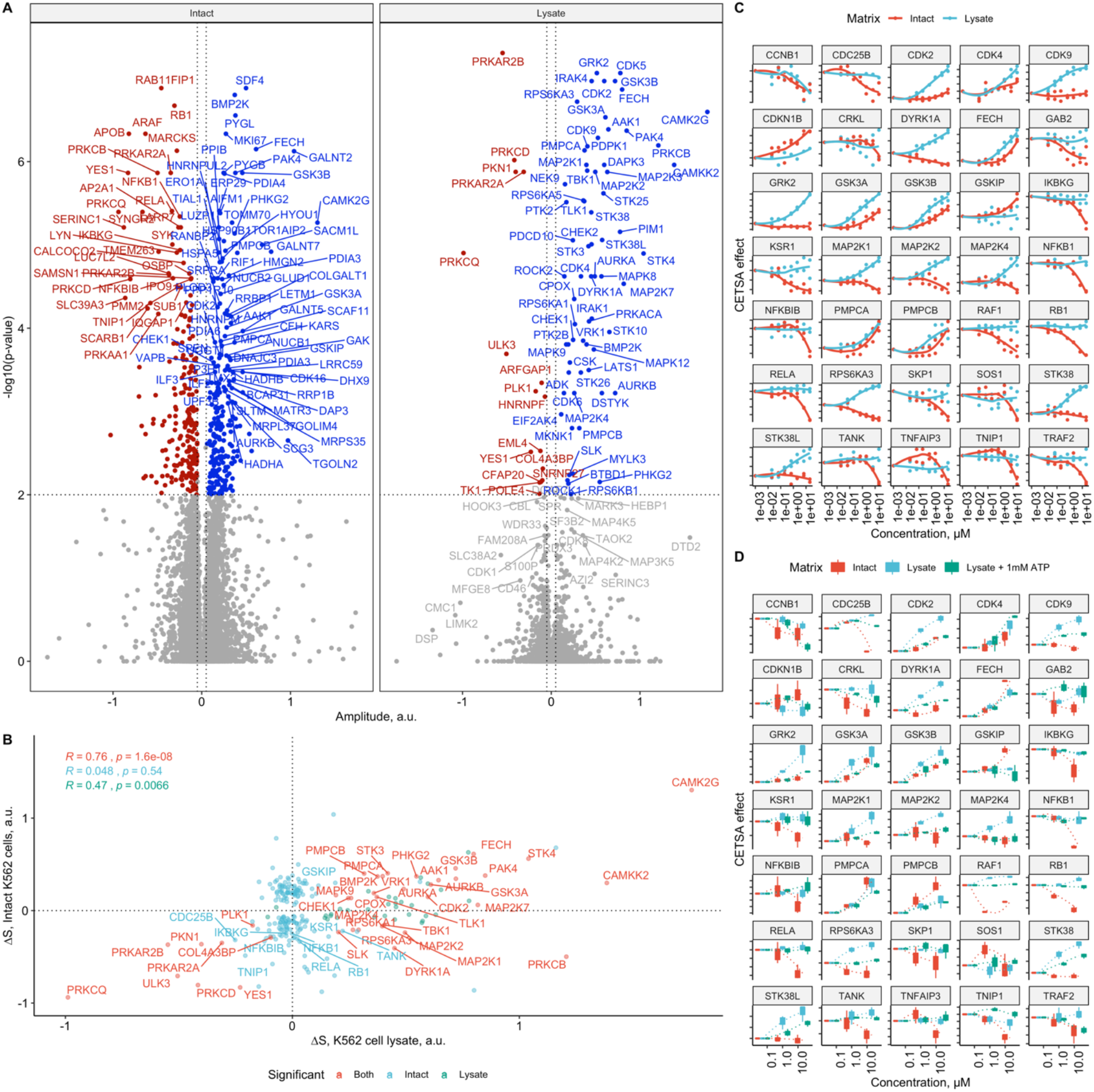
Compressed CETSA MS results for eight concentration point Staurosporine profiling in intact K562 cells and cell lysates. A – volcano plots, B – scatter plot for ΔS amplitudes in both experiment; proteins showing stability changes in both experiments shown in red, lysate-specific in green, and intact only protein hits shown in blue. C – concentration response profiles for selected proteins from eight-concentration point compressed CETSA MS experiment. D – concentration dependent changes in thermal stability of selected proteins in the three-concentration point experiment in intact K562 cells, cells lysate and cell lysate with 1mM ATP added.

Some of the kinases that were thermally stabilized in the lysate format were instead found to be destabilized in the intact cell experiment, which was also observed for many of the non-kinase interacting proteins in intact cells. An example is MAP2K1(MEK1), which was stabilized in lysate but destabilized in intact cells. The MAP2K1 interacting kinases KSR1 and RAF1 were only found to be hits in intact cells and similarly to MAP2K1 they were also destabilized (Figure 3C). The tilt towards thermal destabilization in intact cells suggests that interacting proteins that are likely to reside in complexes are more prone to get destabilized upon ligand binding in contrast to dispersed proteins in a lysate. Support for similar thermal stability of interacting proteins have previously been observed by Savitski et al.^2^, and later also reported by Tan et al.^5^

The realization that there were substantial differences between lysate and intact cell experiment was intriguing. The reason for this difference could be the dispersion of protein complexes in lysates, but it could also be attributed to the decreased concentration of small metabolites, mainly ATP, in the lysates.

In order to test the hypothesis that the difference between intact and lysate results was due to the ATP levels, we performed another lysate experiment in presence of a physiological concentration of ATP (1 mM). We could then detect a decreased sensitivity to heat treatment in the presence of ATP for some kinases, whereas others were unaffected, or their response was shifted from stabilization to destabilization (Figure 3D, Supplementary figure 4).

### Kinases not-shifting upon Staurosporine treatment

Detecting proteins which do change stability upon treatment is a relatively straightforward task, however, confidently assigning proteins as non-shifted proteins is much more challenging. In order to approach this question, we have accumulated data from 14 individual biological replicates of Staurosporine-treated K562 lysates, which allowed us to identify and reliably quantify more than 7500 proteins. Using the power of such number of replicates we have been able to detect 293 proteins with concentration-dependent changes in thermal stability: 169 proteins (117 kinases) stabilized and 63 (11 kinases) destabilized proteins. Surprisingly, increasing the number of replicates from four (the first Compressed experiment above) to fourteen did not change the number of destabilized kinases.

The majority of non-kinase hits identified with relatively low effect size and has at least one phosphorylation site according to their Uniprot annotations. This suggests that thermal stability changes for those proteins happen, not due to the Staurosporine binding, but rather that some level of kinase activity remains in lysed cells, and this activity could be modulated by Staurosporine. However, depth of the dataset and number of replicates allows us to select a subset of kinases which do not change their stability upon Staurosporine treatment. We have mapped the results from this experiment to the kinome tree using KinMap service (http://www.kinhub.org/kinmap/) (Figure 4). Stabilized kinases shown in blue circles, destabilized-in red circles, while those kinases which definitely remained unshifted shown as grey squares. The size of the circles reflects pXC50 values.

**Figure 4.**
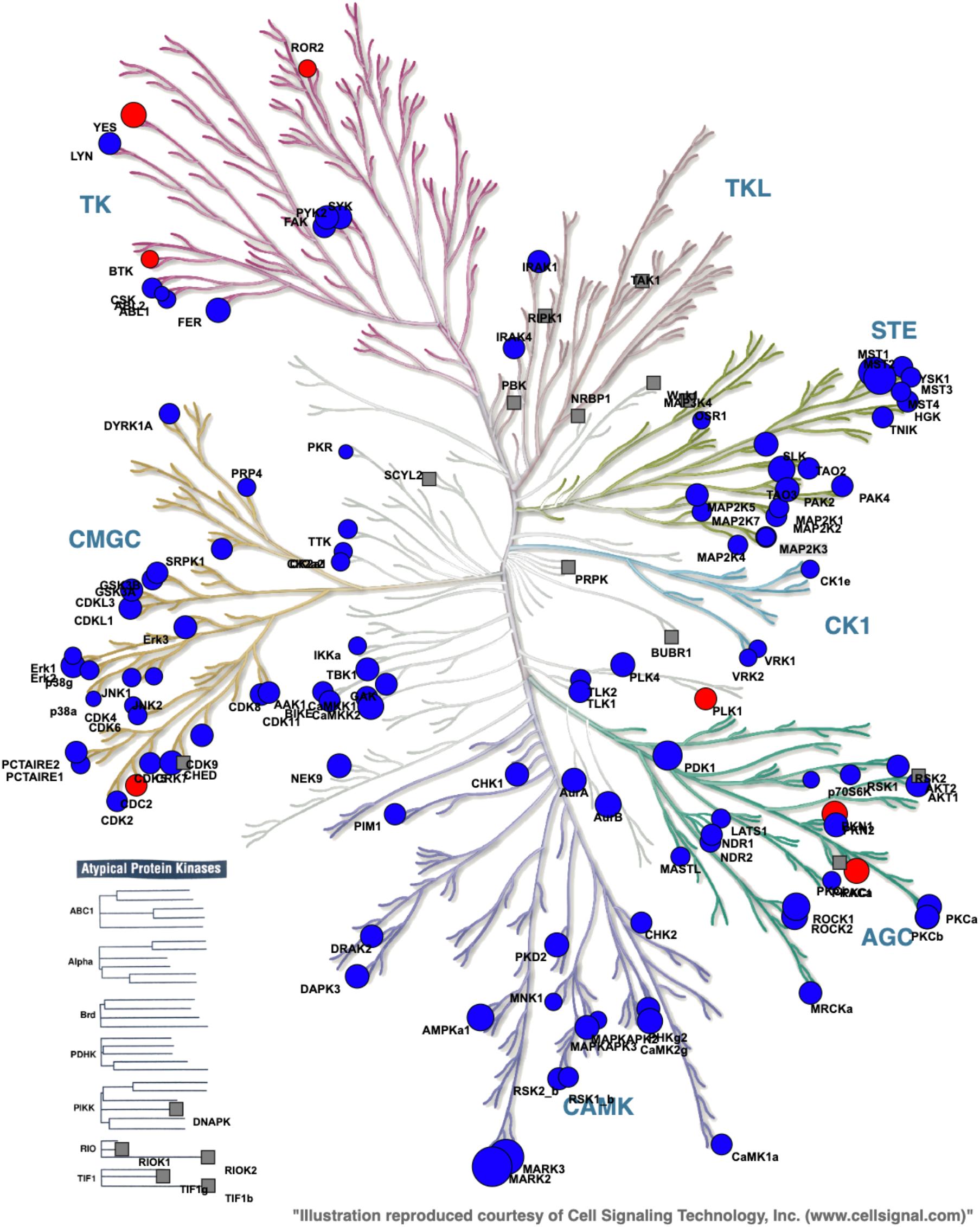
Map of human kinases identified in eight-concentration point compressed CETSA MS experiment performed in K562 cell lysate. Kinases stabilized upon Staurosporine treatment shown in blue, destabilized – in red, circle size is proportional to the extracted pXC50 value. No-shift kinases (with enough evidence) shown as gray squares.

## Conclusions

Since its introduction 2013, CETSA has received much attention as a means to demonstrate physical target engagement of added compound. With the introduction of multiplexed mass-spectrometric read out, researchers are now able to look beyond the direct target engagement measure. CETSA MS reports on basically all changes in the cell that affect protein stability, be it protein-protein interactions, phosphorylation state or sub-cellular protein relocalization induced by the added drug. This way, CETSA MS could offer robust measures of cellular responses, including the generation of biomarkers for treatment efficacy.

In this report we have used Staurosporine, a well-known and promiscuous kinase inhibitor to highlight differences, pros and cons, of different experimental layouts. We compare the use of the qualitative melt curve approach with the quantitative dose-response format, as well as the fusion of the both, the so called 2D-TPP. We have carried out the experiments in both intact cells and lysate to highlight the differences between a system with intact cellular signaling machinery and a system reporting exclusively on direct binding. We find that we generally have a good reproducibility and overlap between the different experimental layouts but as expected, more datapoints give a higher accuracy and better statistical ground to confidently assign proteins as responders or non-responders. Having been available to the wider community for only 6 years, CETSA MS is still in its infancy with regards to finding optimal protocols and experimental layouts. With an ever-increasing understanding of the power (and resolution) of the method, the number of potential applications is likely to grow. In addition, with technological advancements, such as the introduction of novel isobaric labelling chemistry^12^ and LC-MS throughput^13^ it becomes possible to analyze large number of compounds and unravel common or distinct pathways activation.

## Material and methods

K562 cells were obtained from ATCC and were cultured following ATCC protocols. Cell cultures were kept sub-confluent and the cells were given fresh medium one day prior to harvesting. For the experiment, the cells were washed and detached with 1x TrypLE (Gibco, Life Technologies). The detached cells were diluted into Hank’s Balanced Salt solution (HBSS), pelleted by centrifugation (300 g, 3 min), washed with HBSS, pelleted again and re-suspended in CETSA® buffer (20 mM HEPES, 138 mM NaCl, 5 mM KCl, 2 mM CaCl2, 1 mM MgCl2, pH 7.4) constituting the 2x cell suspension. For the lysate experiments, K562 cells were re-suspended in CETSA® buffer and cells were lysed by three rounds of freeze-thawing. The lysate was clarified by centrifugation at 30 000 x g for 20 minutes and used immediately in the CETSA® experiment constituting the 2x lysate. Reagents were purchased from Sigma unless otherwise noted.

### CETSA MS melt curve and five-concentration points compressed experiment

The K562 lysate was divided into ten aliquots (five concentration points in duplicates) and each was mixed with an equal volume of each test condition prepared at 2x final concentration in the experimental buffer (20 mM HEPES, 138 mM NaCl, 5 mM KCl, 2 mM CaCl2, 1 mM MgCl2, pH 7.4). The resulting final concentrations of Staurosporine was 0.02, 0.2, 2, and 20μM; 1% DMSO only was used as control. Incubations were performed for 15 minutes at room temperature with end-over-end rotation.

The treated cell suspensions were each divided into ten aliquots that were all subjected to a heat challenge for 3 minutes, each at a different temperature between 40 and 67°C. After heating, precipitated protein was pelleted by centrifugation at 20 000 g for 20 minutes and supernatants containing soluble fractions were kept for further analysis.

The total protein concentration of the soluble fractions was measured by Lowry assay DC (BioRad). For the melt-curve CETSA MS experiments only samples from highest compound concentration (20μM) and vehicle control (1% DMSO) were taken for the further analysis. Same volume from all individual temperatures was taken. It resulted in forty samples (ten temperatures, treatment vs control, two replicates). For the compressed CETSA experiment same volume from individual temperatures were pulled together for each of the experimental conditions producing 10 individual samples (five concentration points in two replicates). Each of these samples were split in two replicates, resulting in total of 20 samples.

Samples were subjected to reduction and denaturation with tris(2-carboxyethyl)phosphine (TCEP) (Bond-breaker, Thermo Scientific) and RapiGest SF (Waters), followed by alkylation with chloroacetamide. Proteins were digested with Lys-C (Wako Chemicals) for 6 hours and trypsin (Trypsin Gold, Promega) overnight.

After complete digestion had been confirmed, samples were labelled with 10-plex Tandem Mass Tag reagents (TMT10, Thermo Scientific) according to the manufacturer’s protocol. Labelling reactions were quenched by addition of a primary amine buffer and the concentration series from two consecutive temperatures were combined in one TMT10-plex set (rows in Table 1). The labelled samples were subsequently acidified and desalted using polymeric reversed phase (Strata X, Phenomenex). LC-MS grade liquids and low-protein binding tubes were used throughout the purification. Samples were dried using a centrifugal evaporator.

### LC-MS/MS analysis

For each TMT10-plex set, the dried labelled samples were dissolved in 20 mM ammonium hydroxide (pH 10.8) and subjected to reversed-phase high pH fractionation using an Agilent 1260 Bioinert HPLC system (Agilent Technologies) over a 1 × 150 mm C18 column (XBridge Peptide BEH C18, 300 Å, 3.5 μm particle size, Waters Corporation, Milford, USA). Peptide elution was monitored by UV absorbance at 215 nm and fractions were collected every 30 seconds into polypropylene plates. The 60 fractions covering the peptide elution range were evaporated to dryness, ready for LC-MS/MS analysis.

Thirty individual fractions were analyzed by high resolution Q-Exactive HF Orbitrap mass spectrometers (Thermo Scientific) coupled to high performance nano-LC systems (Evosep One, Evosep, Denmark). All samples were injected in duplicate for replicate data collection.

MS/MS data was collected using higher energy collisional dissociation (HCD) and full MS data was collected using a resolution of 120 K with an AGC target of 3e6 over the m/z range 375 to 1500. The top 12 most abundant precursors were isolated using a 1.4 Da isolation window and fragmented at normalized collision energy values of 35. The MS/MS spectra (45 K resolution) were allowed a maximal injection time of 120 ms with an AGC target of 1e5 to avoid coalescence. Dynamic exclusion duration was 30 s.

### Data analysis

Protein identification was performed by database search against 95 607 canonical and isoform human protein sequences in Uniprot (Swissprot and TrEMBL, UP000005640, download date: 2019-02-21) using the Sequest HT algorithm as implemented in the ProteomeDiscoverer 2.2 software package. Data was re-calibrated using the recalibration function in PD2.2. and final search tolerance setting included a mass accuracy of 10 ppm and 50 mDa for precursor and fragment ions, respectively. A maximum of 2 missed cleavage sites was allowed using fully tryptic cleavage enzyme specificity (K\, R\, no P). Dynamic modifications were; oxidation of Met, and deamidation of Asn and Gln. Dynamic modification of protein N-termini by acetylation was also allowed. Carbamidomethylation of Cys, TMT-modification of Lysine and peptide N-termini were set as static modifications.

For protein identification, validation was done at the peptide-spectrum-match (PSM) level using the following acceptance criteria; 1 % FDR determined by Percolator scoring based on Q-value, rank 1 peptides only.

For quantification, a maximum co-isolation of 50 % was allowed. Reporter ion integration was done at 20 ppm tolerance and the integration result was verified by manual inspection to ensure the tolerance setting was applicable. For individual spectra, an average reporter ion signal-to-noise of ≥20 was required. Only unique peptide sequences were used for quantification.

## Supplementary figures

**Supplementary figure 1.**
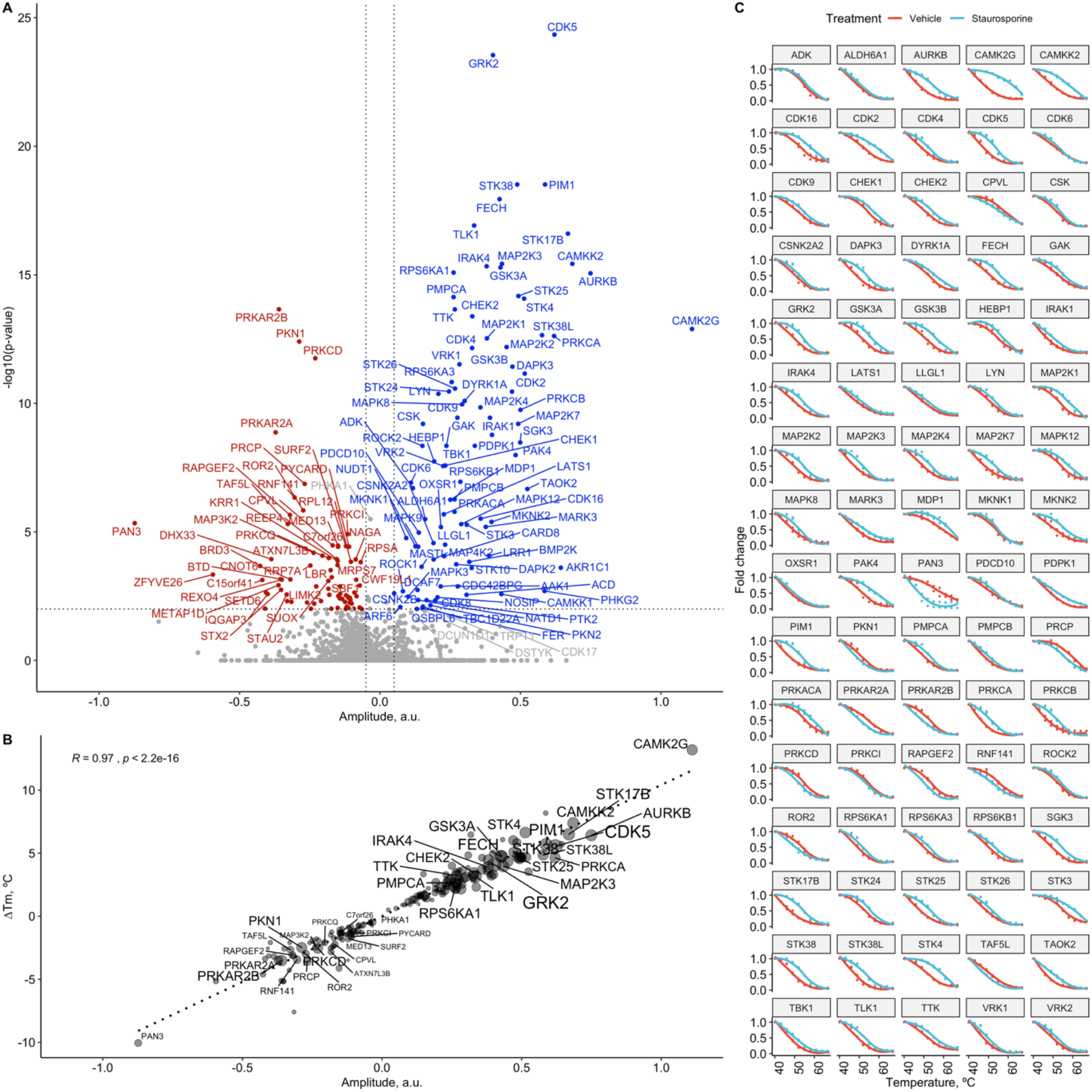
Summary of CETSA MS profiing of Staurosporine in K562 cell lysate performed in a melt-curve format. A – volcano plot with proteins significantly stabilized shown in blue, destabilized – blue, non-significant – gray.B – scatter plot showing correlation between calculated ΔT_M_ and ΔS values. C – melt curves for 80 most significant protein hits.

**Supplementary figure 2.**
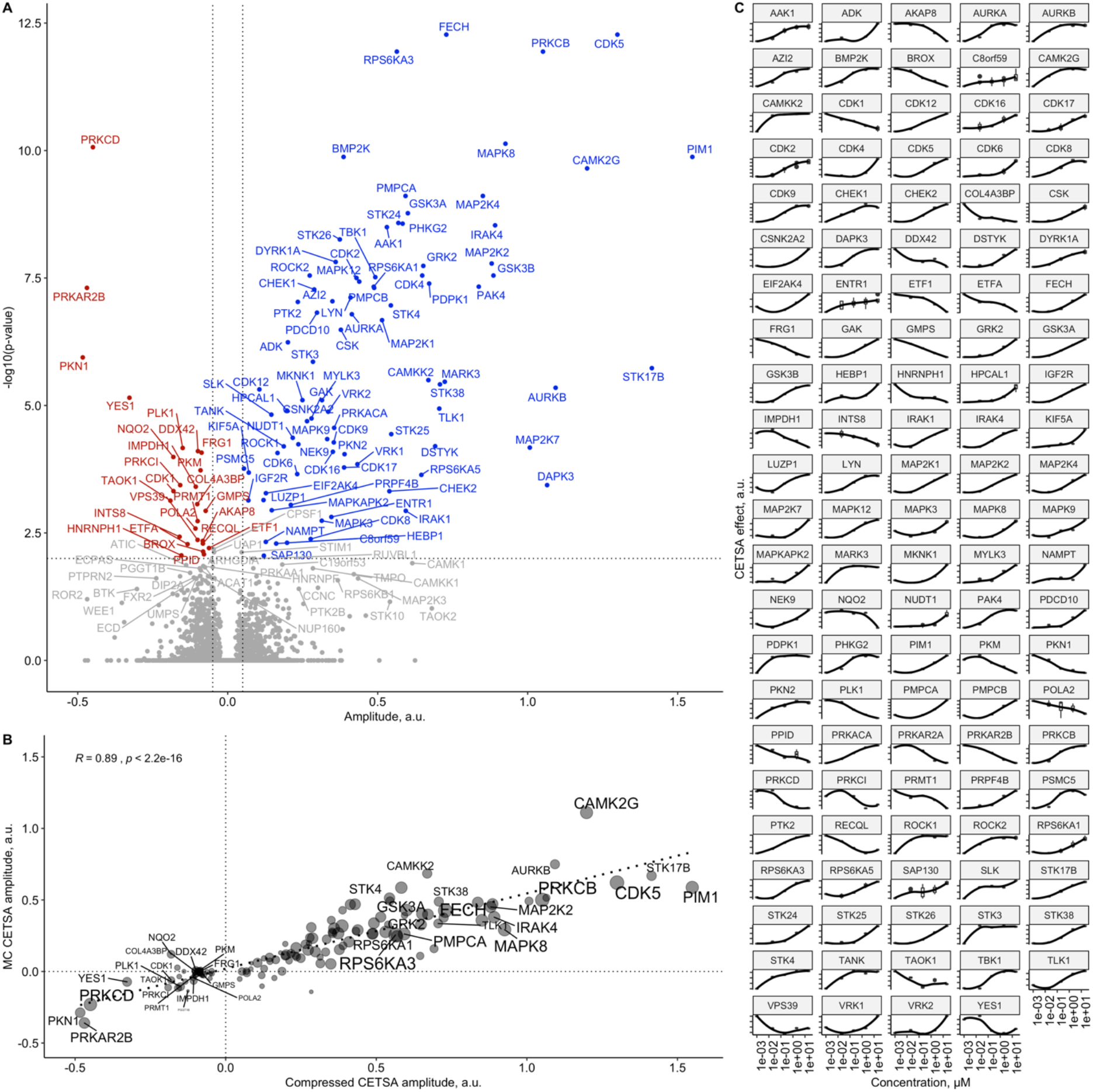
Summary of CETSA MS profiing of Staurosporine in K562 cell lysate performed in five-concentration point compressed format. A – volcano plot with proteins significantly stabilized shown in blue, destabilized – blue, non-significant – gray. B – scatter plot showing correlation between measured ΔS values and ΔS values calculated from melt curve experiment.. C – concentration response profiles for 80 most significant protein hits.

**Supplementary figure 3.**
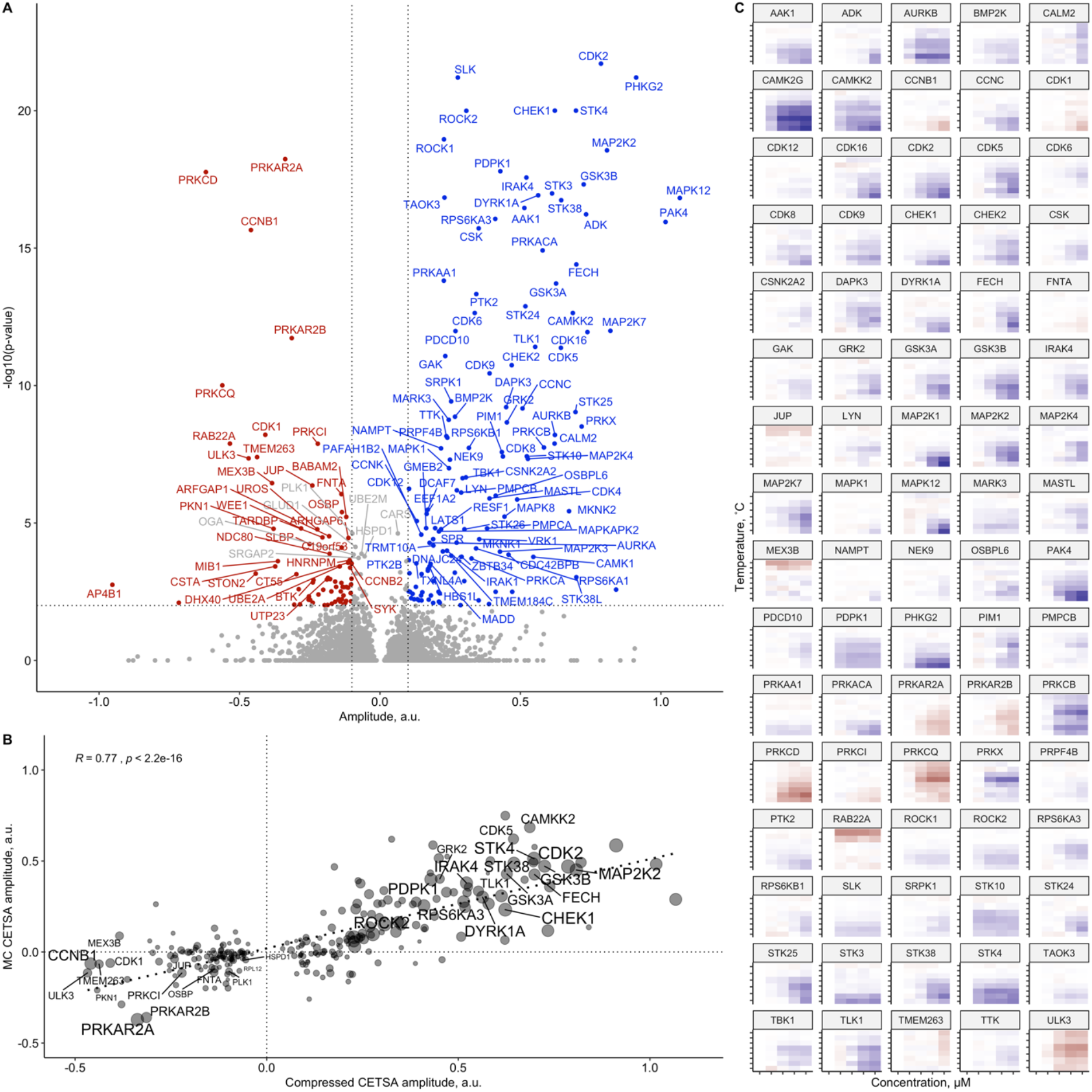
Summary of CETSA MS profiing of Staurosporine in K562 cell lysate performed in five-concentration point two-dimensional (2D) format. A – volcano plot with proteins significantly stabilized shown in blue, destabilized – blue, non-significant – gray. B – scatter plot for measured ΔS value versus ΔS values calculated from melt-curve experiment. C – 2D-heatmaps for 80 most significant protein hits (color-coded foldchanges are shown, with blue means stabilization and red-destabilization).

**Supplementary figure 4.**
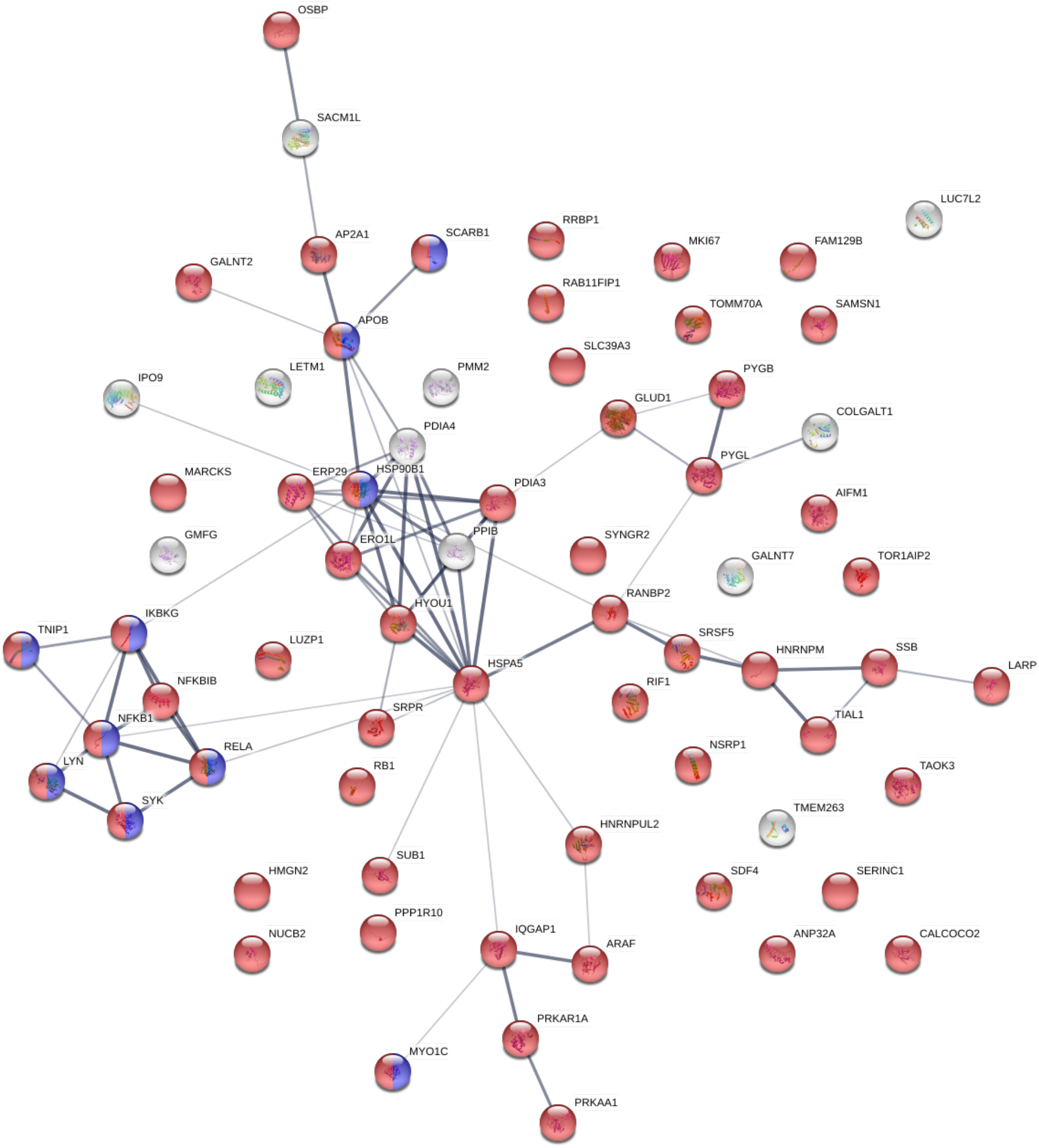
Proteins found to be significantly stabilized upon Staurosporine treatment in intact K562 cells, but not in K562 cell lysate. Phosphoproteins shown in red.

**Supplementary figure 5.**
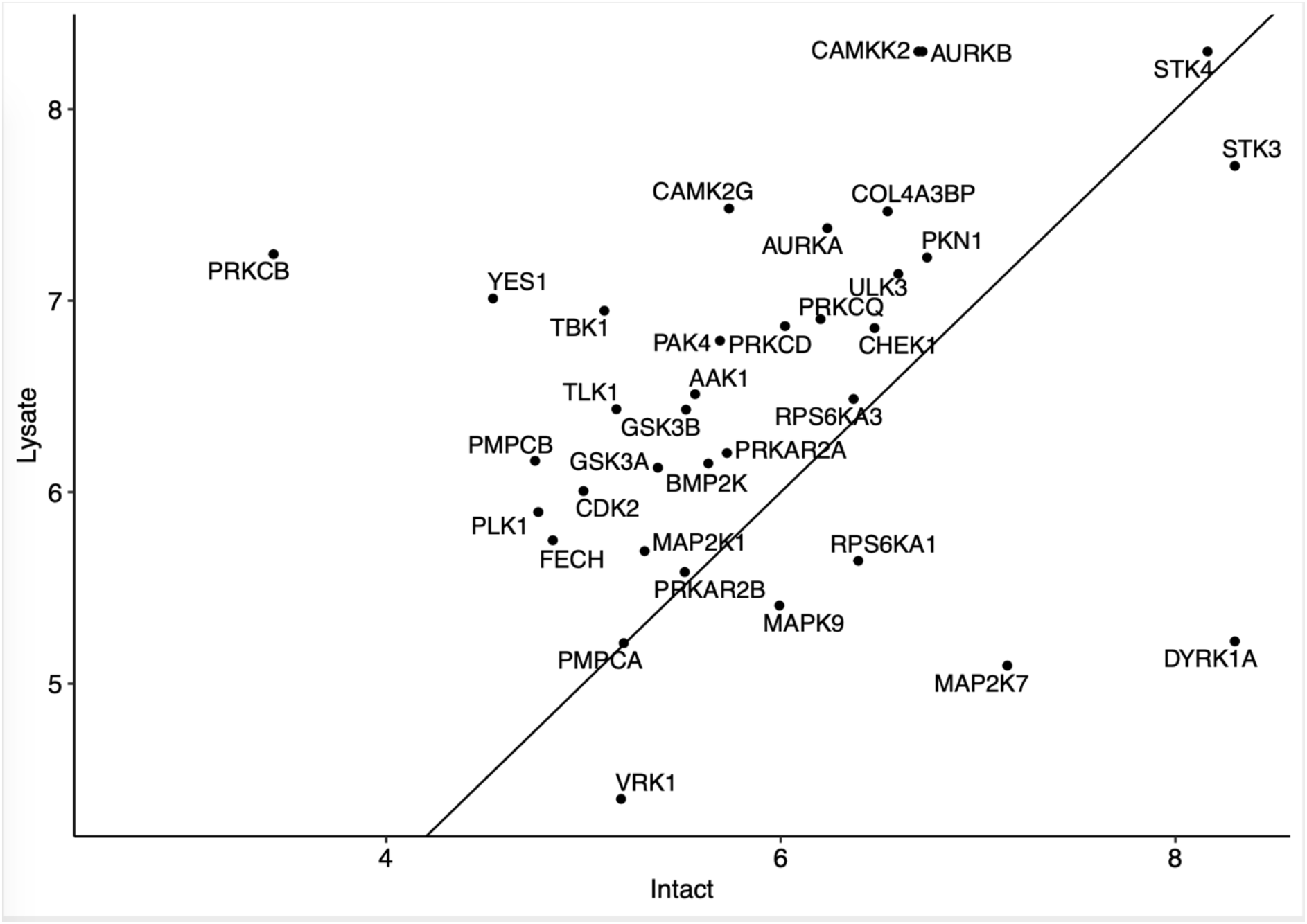
XC50 values from eight-concentration point compressed CETSA MS experiment in intact K562 cells and cell lysates.

